# Nitrosomes: protein language modeling and live-cell imaging reveal condensate-like nitrogenase organization in heterocysts

**DOI:** 10.64898/2026.05.23.727275

**Authors:** James Young, Shengni Tian, Liping Gu, Dillon Nelson, Huilan Zhu, Jaimie Gibbons, Taufiq Nawaz, Ruanbao Zhou

**Author notes:** Correspondence (J.Y.), (R.Z.).

## Abstract

Biological nitrogen fixation in some filamentous cyanobacteria occurs in heterocysts, yet the subcellular organization of nitrogenase in these cells remains poorly defined. Using fluorescence microscopy of living *Anabaena* sp. PCC 7120, we found that a NifH-GFP fusion forms discrete puncta restricted to mature heterocysts, whereas constitutively expressed GFP in vegetative cells and a nifB-linked GFP reporter remained diffuse. To contextualize this phenotype, we built a homology-aware cyanobacterial condensate-prioritization framework across 31,028 proteins from seven proteomes, integrating ESM-2 embeddings, 27 biophysical features, and 44 curated condensate-associated positives. Because direct positives remain limited, we present the model as a ranking resource rather than a calibrated classifier. Under a deliberately conservative nitrogenase-withheld analysis that removed nitrogenase-family labels and homologous clusters from the training data, NifH retained a median rank in the highest-scoring 4.8% of cyanobacterial proteins. catGRANULE 2.0 also assigned high scores to nitrogenase iron proteins, while having only modest rank concordance to the whole atlas. Together, these data identify heterocyst-restricted NifH-GFP puncta as evidence of sub-cellular spatial organization and provide a curated resource for prioritizing cyanobacterial condensate candidates, some of which are also important for nitrogen fixation.

## 1. Introduction

Biomolecular condensates formed by liquid-liquid phase separation (LLPS) organize biochemical processes across all domains of life^1,2^. These non-membranous compartments concentrate specific molecular components, allowing cells to spatially organize metabolism, regulate gene expression, and respond to environmental stress. Condensate biology has been extensively studied in eukaryotes, including stress granules, P-bodies, and the nucleolus. However, the role of LLPS in prokaryotes, and more specifically in nitrogen-fixation, has received relatively less attention.

Evidence shows that bacterial cells use condensate-like organization in multiple processes. PopZ condensates organize chromosome segregation in *Caulobacter crescentus*^*3,4*^, SSB forms DNA repair foci in *Escherichia coli*^*5*^, and RNA degradosomes assemble as liquid-liquid phase-separated bodies in α-proteobacteria^6^. These findings indicate that bacterial cells are not uniformly mixed cytoplasmic compartments and that condensate organization occurs across bacterial lineages.

Cyanobacteria are an excellent system for studying prokaryotic condensate biology. These organisms perform complex metabolic processes including carbon fixation, nitrogen fixation, and photosynthesis that require spatial organization to manage incompatible chemistries and coordinate electron flow. Many cyanobacterial proteins contain intrinsically disordered regions characteristic of condensate drivers. One example, carboxysomes (proteinaceous microcompartments that concentrate RuBisCO for carbon fixation), involve liquid-like condensation steps during biogenesis^7^, indicating connections between shell-based and condensate-based organization.

Identifying condensate-forming proteins in cyanobacteria remains difficult because few proteins have been directly tested for LLPS capacity in vitro, puncta formation in microscopy can be conflated with true phase separation, and true biological negatives are essentially absent. Cyanobacteria also span a wide phylogenetic range, from unicellular species such as *Synechocystis* and *Synechococcus* to filamentous heterocyst-forming organisms such as *Nostoc* and *Anabaena* and marine nitrogen-fixers such as *Trichodesmium*. These features make cyanobacteria biologically compelling but also demand careful label curation, proxy-negative design, and homology-aware evaluation.

Given the above motivations and limitations, we combined fluorescence imaging of NifH-GFP in heterocysts with a cyanobacterial ranking workflow that uses curated condensate evidence, ESM-2 embeddings, hand-crafted biophysical descriptors, proxy-negative labels, and homology-aware cross-validation to prioritize candidate condensate drivers across seven cyanobacterial proteomes totaling 31,028 proteins as outlined in Figure 1 below. Protein language models, particularly ESM-2^8^, numerically represent implicit concepts such as protein function, structure, and evolutionary relationships. PLM embeddings encode information about intrinsic disorder, domain architecture, and evolutionary conservation that relate to condensate propensity without requiring explicit feature engineering for each property.

**Figure 1.**
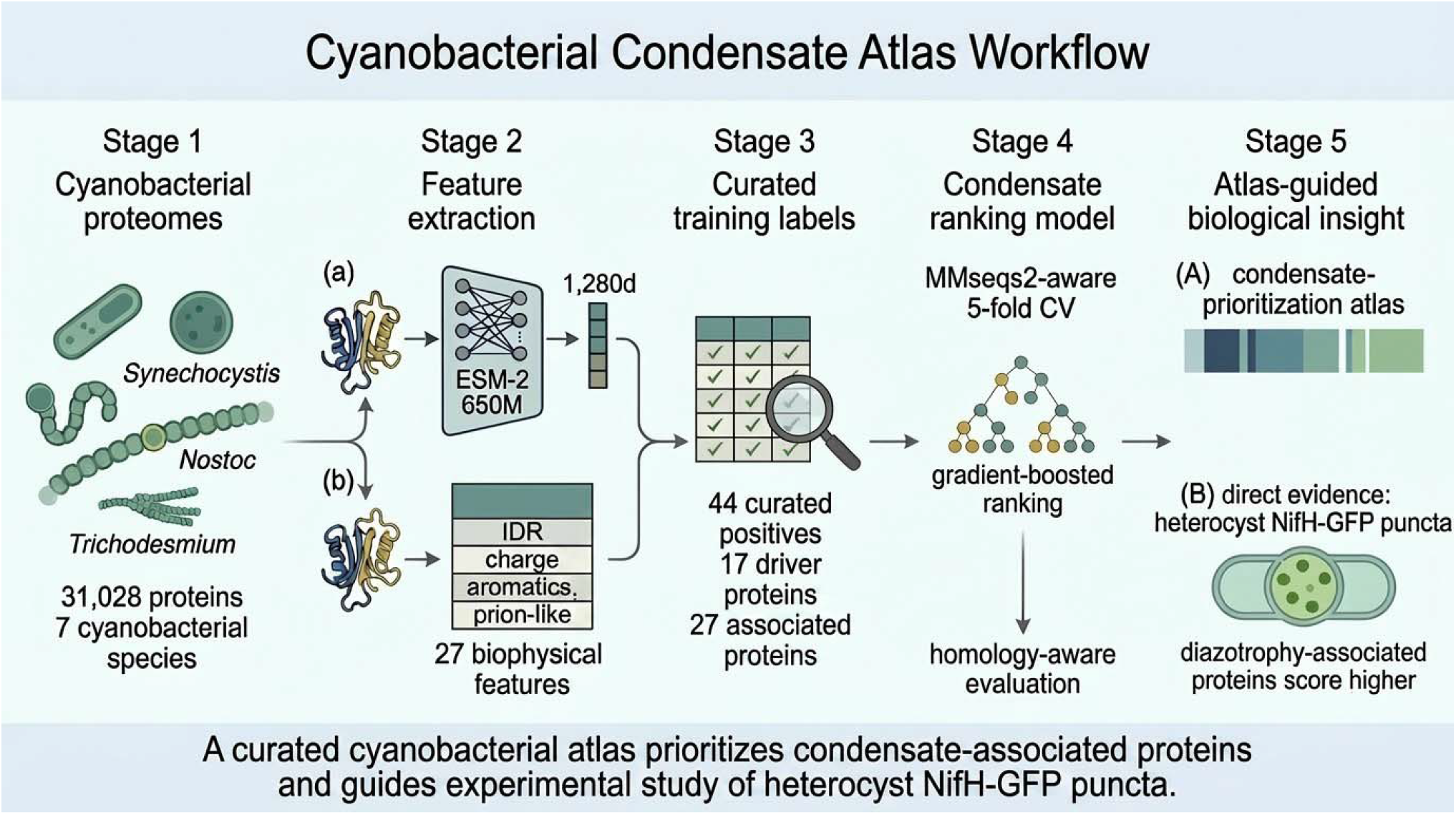
Cyanobacterial condensate-prioritization workflow. Sequences from seven cyanobacterial proteomes (31,028 proteins) are represented by ESM-2 650M embeddings and 27 biophysical features,then scored by a homology-aware gradient-boosted ranking model trained on 44 curated positives (17 driver, 27 associated). The output is a ranking resource that prioritizes condensate-associated proteins and contextualizes heterocyst NifH-GFP puncta imaging.

## 2. Methods

### 2.1 Label Curation

We assembled a curated set of condensate-associated proteins using a four-tier evidence system and performed accession-level identity checks against the final 31,028-protein universe. Rows with mismatched gene-accession pairs were corrected when the intended protein was present in the final universe and otherwise excluded. The corrected label file is provided at data/curated/condensate_labels_corrected.tsv, with explicit copies of the primary direct-curated and homolog-expanded label sets at data/curated/condensate_labels_rescued_direct.tsv and data/curated/condensate_labels_rescued_direct_plus_homologs.tsv. The primary curated file contains 44 training-eligible positives: 2 condensate-evidence Tier 1 proteins, 26 Tier 2 proteins, and 16 Tier 3 proteins. The two condensate-evidence Tier 1 driver anchors are *Escherichia coli* SSB (P0AGE0) and FtsZ (P0A9A6).

For model training, condensate-evidence Tier 1 through Tier 3 rows were eligible as positives when their label_type matched the target, yielding 17 condensate-driver positives and 27 condensate-associated positives in the primary curated atlas. The NifH-GFP imaging result from this study is represented by the current-study NifH1/NifH2 accessions P00457 and O30577 in the primary atlas, but all nitrogenase-family labels and their homologous clusters were removed from the training data in the independence analysis described below. The conservative 23-row set and 53-row homolog-expanded set are retained as sensitivity analyses.

### 2.2 Feature Representation

Each protein was represented by two feature classes:

#### ESM-2 embeddings

We extracted mean-pooled per-residue embeddings from ESM-2 using both 8M parameter (esm2_t6_8M_UR50D, 320 dimensions) and 650M parameter (esm2_t33_650M_UR50D, 1280 dimensions) variants. Sequences exceeding 1,022 residues were truncated. The final atlas, discussed in this manuscript, specifically uses 650M embeddings for all 31,028 proteins in the unified universe. Principal component analysis (PCA) with 95% variance retention was applied to the 650M embedding dimensions within each training fold, and then the 27 sequence features were appended; fitting PCA inside the fold prevents information leakage from the held-out proteins.

#### Biophysical sequence descriptors (27 features)

Intrinsically disordered region (IDR) fraction (estimated as the fraction of disorder-promoting residues A, Q, S, G, P, E, K, R, D), net charge per residue, mean Kyte-Doolittle hydropathy, aromatic residue fraction (F+Y+W), molecular weight, individual amino acid fractions (20 features), prion-like domain score (maximum Q+N enrichment in an 80-residue sliding window), and sequence length.

The unreduced combined representation contains 1,307 numeric dimensions per protein (1,280 ESM-2 650M dimensions plus 27 sequence descriptors). Production cross-validation and final scoring use PCA-reduced ESM dimensions plus the 27 uncompressed sequence descriptors.

### 2.3 Proxy-Negative Strategies

We employed three proxy-negative strategies, each representing different assumptions about what constitutes a non-condensate-driver. The random-proteome strategy sampled non-positive proteins from the candidate universe and assumes that most proteins are not condensate drivers. The low-IDR structured strategy sampled proteins with IDR fraction below 0.40, which are less likely to drive IDR-mediated phase separation. The phylogenetically matched strategy sampled non-positive proteins from the same organism as each positive with IDR fraction below 0.25, producing a phylogenetically balanced negative set. For each strategy, negatives were sampled at a 5:1 ratio relative to positives. Primary reported tables use random-background proxy negatives. Low-IDR and phylogenetically matched negatives were explored as sensitivity checks but are not the basis of the headline manuscript tables.

### 2.4 Model and Evaluation

We trained a gradient-boosted classifier (scikit-learn GradientBoostingClassifier, 300 trees, max depth 5, learning rate 0.05, subsample 0.8, random_state set to the active seed) to discriminate positives from proxy negatives. The model outputs an uncalibrated target-specific ranking score, not a calibrated physical probability of phase separation.

Evaluation used 5-fold cross-validation. Unless otherwise stated, all headline AUROC numbers are reported under homology-aware splits: we clustered all 31,028 sequences with MMseqs2 easy-cluster at 30% sequence identity and 80% coverage, producing 13,936 clusters, then assigned whole clusters to folds so that homologous pairs do not straddle the train/test boundary. For every reciprocal-best-hit (seed_id, cyanobacterial_id) pair used in the RBH label-expansion step, we union-find merge the cluster containing the seed and the cluster containing the cyanobacteria hit, forcing both proteins into the same fold. This blocks the specific leakage introduced by RBH expansion. Principal component analysis is fit within each training fold only. We also report the random-split AUROC alongside the homology-aware number for comparison so readers can quantify the credibility adjustment.

Multi-seed stability is reported: we retrain and re-split for each seed, and report the mean and standard deviation of fold-averaged AUROC across seeds where available. We report AUROC, AUPRC, and precision/recall at k for each proxy-negative strategy and each feature subset. After cross-validation, the primary curated atlas model was trained on all eligible labeled data and used to score the entire candidate universe.

To test whether nitrogenase-family scores were independent of label or homolog recovery, we ran a nitrogenase-withheld sensitivity analysis. The withheld set was built from MMseqs2 clusters anchored on NifH, NifD, NifK, NifB, NifE, and NifN family members, plus a protein-name sweep for nitrogenase and nif-family annotations. These nitrogenase-family labels and their homologous clusters were removed from the training data: they were excluded from positives and from proxy negatives before model fitting. The model was then trained on the remaining labels and used to score the withheld nitrogenase-family proteins across seeds. This analysis was repeated for the conservative 23-row, primary 44-row, and homolog-expanded 53-row label sets.

Bootstrap enrichment intervals in Section 3.5 use 2,000 row-level resamples with replacement from the full 31,028-protein atlas universe at the prespecified driver-score thresholds 0.5, 0.7, and 0.9 (seed = 42). Fisher exact p-values are one-sided and Benjamini-Hochberg adjusted across the three thresholds for each score column. Because the threshold odds ratios are label-set sensitive, Mann-Whitney U tests provide the primary continuous score-shift evidence.

### 2.5 Candidate Universe

The candidate universe comprised 31,028 proteins from 7 cyanobacterial species plus a small panel of well-characterized bacterial seed proteins used for label transfer (and excluded from all cyanobacterial-only rank reporting):

| Organism | Proteins | Median length (aa) |
| --- | --- | --- |
| *Nostoc* sp. PCC 7120 | 6,709 | 271 |
| *Nostoc punctiforme* PCC 73102 | 6,610 | 270 |
| *Microcystis aeruginosa* NIES-843 | 5,990 | 185 |
| *Synechocystis* sp. PCC 6803 | 3,509 | 263 |
| *Trichodesmium erythraeum* IMS101 | 2,889 | 234 |
| *Synechococcus elongatus* PCC 7942 | 2,863 | 253 |
| *Thermosynechococcus elongatus* BP-1 | 2,451 | 273 |
| bacterial seeds (5 UniProt + 2 NCBI) | 7 | — |

*Nostoc* sp. PCC 7120 proteins were obtained from NCBI RefSeq GCF_000009705.1 (WP_ accessions) because the UniProt proteome UP000002483 release at the time gathered only 786 curated entries; the NCBI RefSeq assembly provides the complete ∼6,709-protein proteome. Proteome sequences for the other six species were obtained from UniProt (release 2026_01). All numerical results reported here are computed against the unified 31,028-protein universe.

### 2.6 Fluorescence microscopy of NifH-GFP in living cells

#### Construction of the cargo plasmids

Plasmids pZR1198 (P*psbA1*-GFP, GFP-only control), pZR2094 (PnifB-nifB-gfp, nifB-linked GFP reporter/comparator), and pZR2113-C2 (P*nifH2*-NifH-GFP fusion) were constructed as summarized in Table S1, Table S2, and Figures S5–S7. All cloned DNA fragments were PCR-amplified with Phusion High-Fidelity DNA Polymerase (NEB) from *Anabaena* sp. PCC 7120 (referred to as *Anabaena* 7120) genomic DNA or related plasmid DNA using the oligos listed in Table S2. PCR products were cloned into the pCR®2.1-TOPO® vector (TOPO TA Cloning® kit, Invitrogen) and confirmed error-free by Sanger sequencing. After digestion with the appropriate restriction endonucleases (NEB), the cloned target fragments were sub-cloned into the pDU1-derived shuttle vectors pSMC232^9^ and pAM1956^10^.

#### Conjugal transfer of cargo plasmids from *E. coli* to *Anabaena*

Cargo plasmids pre-methylated by the methylases remained intact in *mcrBC E. coli* strains (HB101 and NEB10β were used here). *E. coli* HB101 carrying pRL623 plus the conjugal helper pRL443 was mated with *E. coli* NEB10β harboring each cargo plasmid in turn (pZR1198, pZR2094, or pZR2113-C2). The resulting NEB10β strain bearing each cargo plasmid plus pRL443 and pRL623 was then mated with *Anabaena* 7120 to produce, respectively, strains A1198, A2094, and A2113-C2. The conjugal transfer protocol followed ^11,12^.

#### Bacterial strains and growth conditions

Bacterial strains used in this study are listed in Table S1. *Anabaena* 7120 and its genetic derivatives were grown in AA/8 liquid medium with or without nitrate at 30 °C and 100 rpm in an Innova-44R shaker (New Brunswick) under continuous white-light illumination (∼75 µE m^−2^ s^−1^). Liquid cultures of mutant strains were supplemented with 25 µg mL^−1^ neomycin (Nm); 50 µg mL^−1^ Nm was added to agar-solidified AA + N (nitrate-containing) plates for exconjugant selection. *E. coli* strains were grown as described previously^13^.

#### Fluorescence microscopic analysis

*Anabaena* 7120 and its genetic derivatives were grown under continuous light in AA/8 medium with neomycin selection. AA/8 medium was used without combined nitrogen ^14^. After 3–6 days of growth, cultures were imaged on an Olympus BX53 upright wide-field epifluorescence microscope equipped with a digital camera. Brightfield images were acquired in matched fields of view, and GFP fluorescence images were acquired using an SAP GFP filter (U-N31043, Olympus) selected to suppress *Anabaena* autofluorescence. Single-cell three-dimensional structure of one mature heterocyst was reconstructed from a confocal z-stack (Figure 2c; Movie S1). Images were assembled into Figure 2 with linear display scaling; no segmentation thresholding was used for the qualitative puncta call. FRAP, 1,6-hexanediol-sensitivity, temperature-shift, and oxygen-biosensor experiments were not performed in this study; the complete planned experimental panel is described in Limitations Section 4.4 and Future directions Section 4.5.

**Figure 2.**
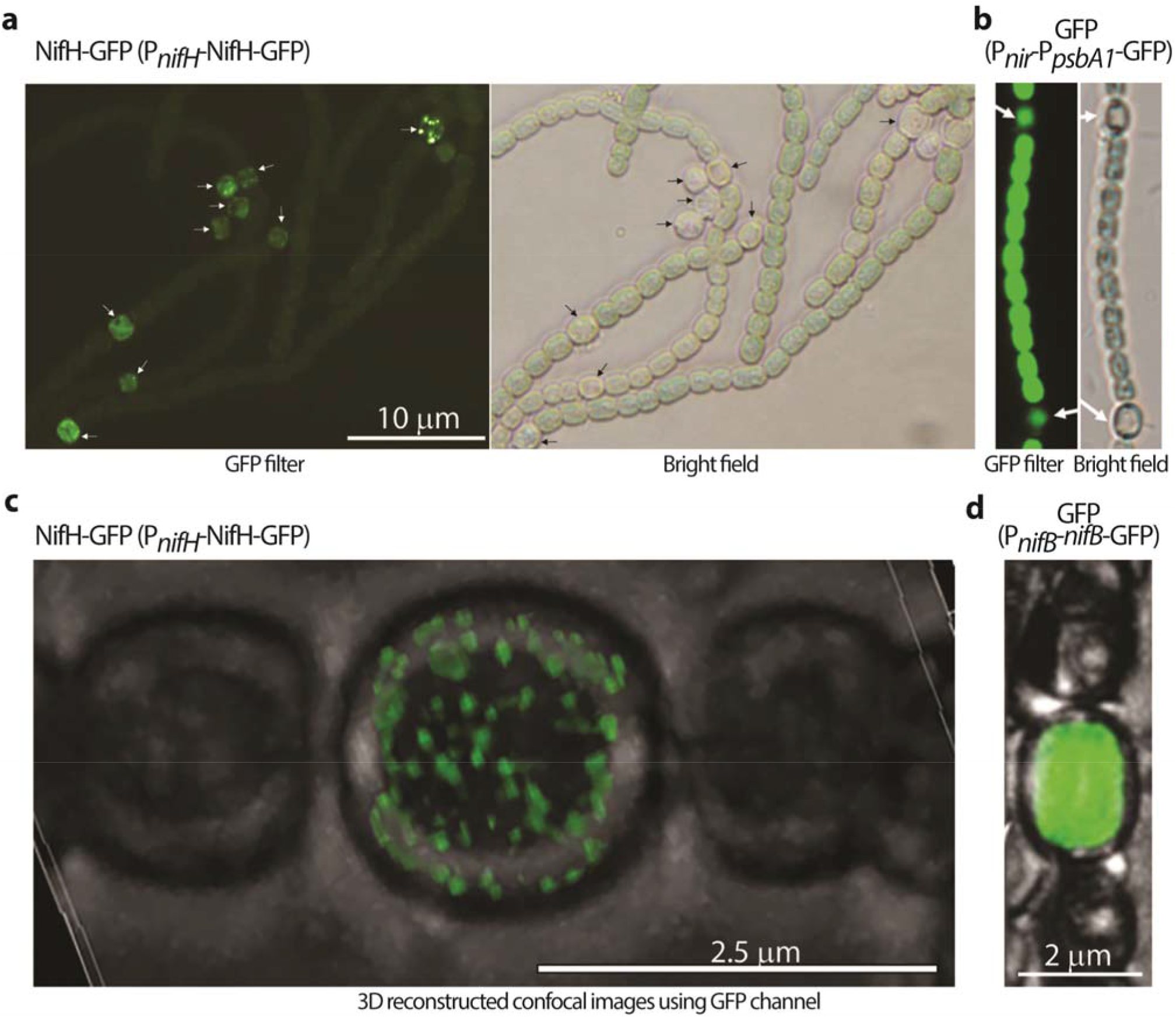
NifH-GFP chimeric protein forms discrete puncta in *Anabaena* (*Nostoc*) sp. PCC 7120 heterocysts. **(a)** Wide-field GFP fluorescence (left) and matched brightfield (right) of filaments of strain A2113-C2 (pZR2113-C2, P_nifH2_-NifH-GFP, NifH is translationally fused to GFP; Tables S1, S2) after six days of growth under continuous light in AA/8 medium with neomycin selection. Discrete spherical puncta form exclusively at heterocyst positions; vegetative cells along the same filament show only background signal. **(b)** Negative control: GFP alone expressed by strain A1198 (pZR1198, P_nir_-P_psbA1_-GFP) shows even diffuse fluorescence in both vegetative cells and heterocysts, with no puncta. **(c)** Higher-magnification 3D-reconstructed confocal image (GFP filter) of one mature heterocyst from strain A2113-C2, resolving discrete puncta filling the cell volume. The corresponding z-stack rotation is provided as Movie S1. **(d)** *nif*B-linked GFP reporter/comparator: strain A2094 carries pZR2094 (P*nif*B-*nif*B-*gfp*); GFP fluorescence was diffuse throughout heterocysts, with no puncta. Imaging in (a, b) at 100× oil; Scale bars: 10 um (a), 2.5 um (c), and 2 um (d). Cultures in panels a–d were grown under matched conditions. AA/8 is a widely used combined-nitrogen-free medium when nitrate is excluded^14^.

**Figure 3.**
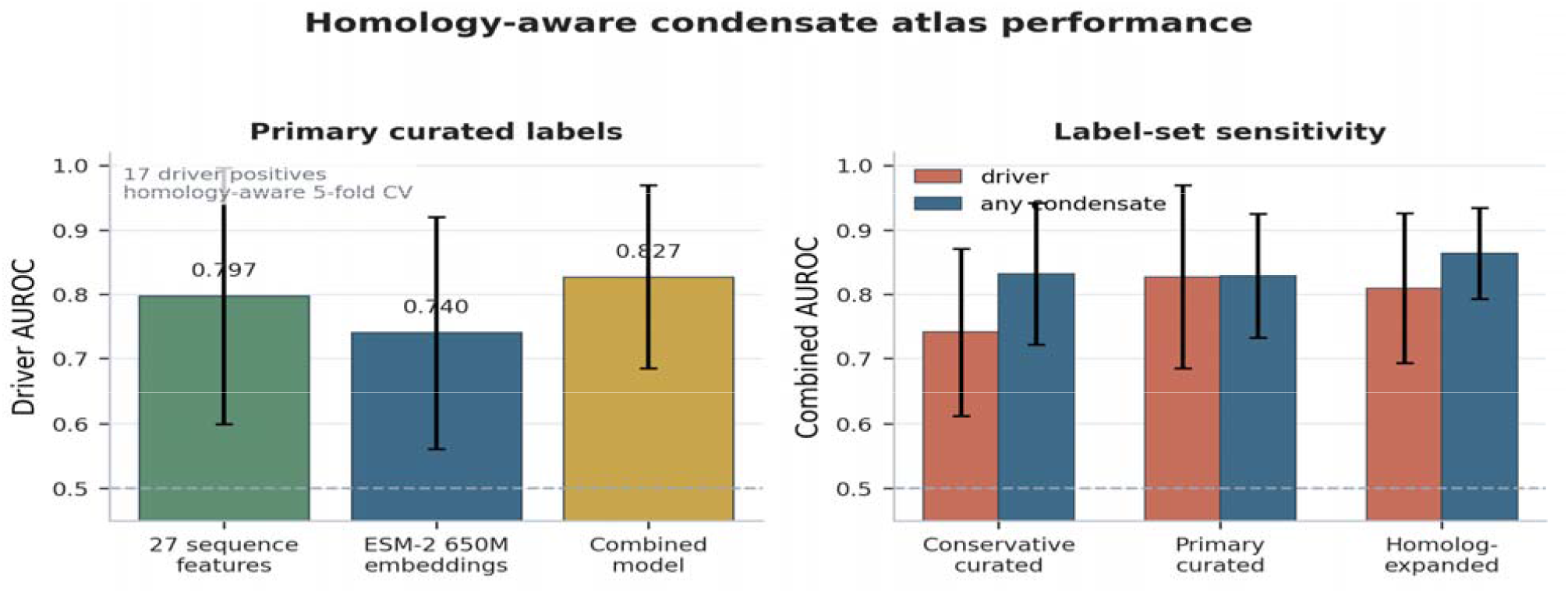
Homology-aware condensate atlas performance. **(A)** Driver AUROC under 5-fold homology-aware cross-validation for the primary curated label set (17 driver positives), comparing 27 biophysical sequence features, ESM-2 650M embeddings with PCA, and the combined representation. **(B)** Combined-model AUROC for driver and any-condensate targets across conservative curated, primary curated, and homolog-expanded label sets. Error bars show across-fold standard deviation.

## 3. Results

### 3.1 Cross-Validation Performance

The accession-aware curation produced a primary 44-positive label set that restores cyanobacterial carboxysome, Rubisco, CcaA, and phycocyanin-associated biology while retaining a conservative 23-positive set for sensitivity analysis. The primary atlas training set contains 17 condensate-driver positives and 27 condensate-associated positives.

Homology-aware cross-validation of the primary curated set gave the following seed-42 values.

| Target | N+ | 27 biophysics only | ESM-2 650M (PCA) only | Combined |
| --- | --- | --- | --- | --- |
| Driver | 17 | 0.797 +/- 0.199 | 0.740 +/- 0.180 | 0.827 +/- 0.142 |
| Any-condensate | 44 | not run for this label set | not run for this label set | 0.829 +/- 0.096 |

Feature ablation showed that the 27 hand-crafted biophysical features carry much of the driver signal in this dataset, with the combined model giving the highest driver AUROC point estimate and ESM-only features performing weaker. Across label-set sensitivity analyses, the combined driver AUROC was 0.741 +/-0.130 in the conservative 23-positive set, 0.827 +/-0.142 in the primary 44-positive set, and 0.809 +/-0.117 in the homolog-expanded 53-positive set. These overlapping intervals argue for a useful but still small-label ranking model rather than a precise performance estimate.

We ran three published sequence-based LLPS predictors trained largely on eukaryotic data (catGRANULE 2.0, PSAP, and LLPhyScore) on 521 cyanobacterial proteins on a Linux HPC (RHEL 9, Python 3.9). This benchmark was computed on the conservative cleaned labeled-plus-background subset, containing 13 driver positives, 21 any-condensate positives, and 500 randomly sampled cyanobacterial background proteins. catGRANULE 2.0 achieved AUROC 0.806 on driver labels and 0.868 on any-condensate labels; PSAP achieved 0.606 and 0.635. LLPhyScore was near chance on driver labels (0.512) and below chance in its published direction for any-condensate labels (0.454; orientation-flipped AUROC 0.546 is shown only as discriminative-capacity context in Supplementary Figures S3 and S4). Same-subset atlas AUROCs are not cited from this benchmark because the labeled subset overlaps the atlas training set; instead, Supplementary Figures S3 and S4 show the external predictors with the homology-aware atlas CV values as reference context. FuzDrop could not be evaluated because it requires ESpritz disorder predictions as input, and the ESpritz precompiled binaries segfault on RHEL 9 / glibc 2.34 while the MobiDB-lite pip alternative requires Python >= 3.12 that was not available on the HPC. Full per-protein scores are provided at results/tables/external_benchmark/, and the conservative benchmark summary is provided at results/tables/sensitivity/external_benchmark_hpc_review.json.

Per-protein concordance showed that the external predictors and the cyanobacterial atlas were complementary rather than redundant. catGRANULE 2.0 and the atlas had similar label-level discrimination but only modest rank concordance across the 521 benchmark proteins (Spearman rho = 0.21 against the driver score and 0.26 against the any-condensate score; top-decile overlaps of 19/52 and 20/52 proteins, compared with 5.2 expected by chance). catGRANULE assigned high scores to heterologous NifH entries, whereas the nitrogenase-withheld atlas deliberately made the Nif-family test harder by removing nitrogenase-family labels and their homologous clusters from the training data before scoring current-study NifH. This convergence from a generic LLPS predictor and a cyanobacterial, homology-aware prioritization model supports NifH as a strong candidate for condensate-mechanism testing while leaving direct LLPS validation unresolved.

Low-IDR negative strategies yield excellent separation (AUROC approaching 1.0) for all models under random splits, reflecting the strong disorder signal in positives rather than genuine ranking quality; we therefore use the random-proteome proxy negatives with homology-aware folds as the primary model-performance estimate.

### 3.2 NifH-GFP forms discrete cytoplasmic puncta in heterocysts

We compared NifH-GFP localization by fluorescence microscopy of living filaments carrying three constructs in *Anabaena* (*Nostoc*) sp. PCC 7120: the NifH-GFP fusion (pZR2113-C2, *P*_*nifH2*_-NifH-GFP, strain A2113-C2), a constitutive GFP control (pZR1198, *P*_*nir*_*-P*_*psbA1*_-GFP, strain A1198), and a heterocyst-expressed GFP comparator (pZR2094, *PnifB*-*nifB*-GFP, strain A2094). Living filaments were imaged after six days of growth in AA/8 medium with neomycin selection (Figure 2). NifH-GFP is located to discrete, spherical, cytoplasmic puncta exclusively in heterocysts (Figure 2a, arrowed); vegetative cells along the filament showed diffused signal. By contrast, GFP alone driven from the constitutive P_nir_-*P*_*psbA1*_ dual promoter (Figure 2b) was evenly diffused throughout both heterocysts and vegetative cells, and GFP fluorescence from the pZR2094 nifB-linked reporter was diffuse throughout heterocysts (Figure 2d). Neither comparator produced puncta. Confocal microscopy of a single mature heterocyst (Figure 2c, Movie S1) resolved spatially distinct puncta filling the heterocyst volume, an order of magnitude more discrete foci than visible in wide-field imaging.

These structures show features compatible with condensate-like organization, including round shape, multiple resolvable occurences per cell, and absence from the tested fluorescent comparators. Morphology alone does not establish liquid-liquid phase separation, composition, function, or membrane independence. We therefore describe the observed structures operationally as NifH-GFP puncta.

The controls reduce, but do not remove, fluorescent-tag artifact concerns. Weak GFP-family oligomerization and fusion context can alter protein localization, clustering, and condensate behavior^18-21^. In the present imaging framework, however, constitutively expressed GFP in vegetative cells and heterocyst-expressed GFP remained diffused while NifH-GFP formed heterocyst-restricted puncta. This argues against a universal “GFP makes puncta” explanation and supports a cargo-dependent NifH-GFP phenotype, while still requiring endogenous tagging, orthogonal fluorophores, expression-level controls, FRAP, perturbation, and ultrastructural validation.

In the primary atlas, NifH-family proteins rank highly when nitrogenase-family labels are allowed in training. The nitrogenase-withheld sensitivity model gives the more appropriate independence test. Across five seeds in the cyano-only universe, current-study NifH1 (P00457) ranked at median 1,497 (IQR [1,363, 8,600]; range 1,273-10,152), corresponding to the highest-scoring 4.8% of cyanobacterial proteins. Current-study NifH2 (O30577) ranked at median 4,429 (IQR [2,727, 4,892]; range 593-8,877), corresponding to the highest-scoring 14.3%. The RefSeq NifH entries tracked these UniProt entries closely. The ChlL paralog Q8YM62 was not counted as a NifH label. Thus, the imaging result should be read as direct experimental evidence for heterocyst-restricted NifH-GFP puncta, with the atlas providing rank-based prioritization rather than a stand-alone nitrogenase-independent nomination.

Several limitations constrain this interpretation. The constitutive GFP control and NifB-GFP comparator (Figure 2b, d) argue against non-specific GFP aggregation as the only explanation, but FRAP has not yet been performed, so mobile fraction and recovery half-time cannot be used to distinguish liquid-like dynamics from static aggregates. Perturbation tests with 1,6-hexanediol, an isogenic PopZ-GFP positive control in the same strain, and temperature-shift dynamics also remain outstanding. Local oxygen tension within NifH-GFP puncta has not been measured, so the proposal that these puncta provide an O2-reduced microenvironment for oxygen-sensitive nitrogenase remains a hypothesis. These controls define the priority follow-up experiments.

What the present data show is that NifH-GFP produces punctate, heterocyst-restricted structures that are absent from the tested fluorescent comparators, and that the nitrogenase-withheld atlas places current-study NifH1 and NifH2 at median cyano-only ranks of 1,497 and 4,429, corresponding to the highest-scoring 4.8% and 14.3% of cyanobacterial proteins.

### 3.3 Ranked Candidate Atlas

Among non-training, non-nitrogenase-family proteins, the primary nitrogenase-withheld cyano-only shortlist is led by interpretable families connected to bacterial spatial organization, including carboxysome assembly proteins, cell-division proteins, DNA repair and single-stranded DNA-binding proteins, Dps/ferritin stress proteins, RNA-binding proteins, and selected electron-transfer proteins. Full cyano-only ranks are shown:

| Cyano driver rank | UniProt | Gene | Protein name | Organism |
| --- | --- | --- | --- | --- |
| 3 | B0JVM0 | *ccmM* | Carboxysome assembly protein CcmM | *Microcystis aeruginosa* NIES-843 |
| 5 | O85785 | *ftsZ* | Cell division protein FtsZ | *Synechococcus elongatus* 7942 |
| 22 | B2J837 | Npun_R5701 | Ferritin/Dps family protein | *Nostoc punctiforme* PCC 73102 |
| 26 | P0A4K0 | *ssb1* | Single-stranded DNA-binding protein 1 | *Nostoc* sp. PCC 7120 |
| 31 | B0JV95 | *recA* | Recombinase A | *Microcystis aeruginosa* NIES-843 |
| 43 | P0A3Y0 | *petJ* | Cytochrome c6 | *Synechococcus elongatus* 7942 |
| 65 | Q44554 | *rbpB* | Putative RNA-binding protein RbpB | *Nostoc* sp. PCC 7120 |

Nitrogenase-family entries are not framed as novel computational nominations because their standard-atlas scores are sensitive to whether nitrogenase-family labels are allowed in training. Several high-ranked entries are close to training families or broad stress-response systems, so their ranking should be used as a prioritization cue rather than as evidence of condensate formation. The renewed prominence of CcmM and other carboxysome-associated biology is notable because it arises after accessions were corrected rather than from stale mismatched rows.

The any-condensate rankings similarly emphasize carboxysome-associated proteins, chaperones, translation machinery, DNA repair proteins, and a smaller set of electron-transfer or signaling proteins. These categories are consistent with known bacterial condensate biology because carboxysome biogenesis involves condensation-like assembly, chaperones and ribosomal proteins partition into bacterial condensates, and RecA forms condensate-like foci during DNA repair.

### 3.4 Feature Importance

The nitrogenase-withheld feature-ablation analysis supports the same conservative reading (Supplementary Figure S2). For seed 42, the sequence-feature model ranked NifH1/NifH2 higher than the ESM-only model, whereas NifK showed the opposite pattern and ranked strongest under the ESM-only model. Across five combined-model seeds, NifH1, NifK, and NifB had median ranks within the top 10% of the cyanobacterial universe, while NifH2, NifE, and NifN were weaker or more variable. This mixed behavior argues against a simple pathway-memorization explanation and supports treating the atlas as a hypothesis-generation tool rather than as a mechanistic classifier.

### 3.5 Nitrogen Fixation Proteins Are Enriched for Condensate Propensity

To assess the biological relevance of our condensate predictions, we cross-referenced the atlas with the diazotrophy-conservation family list deposited with this manuscript (Table S4). Proteins from families enriched for conservation across diazotrophic cyanobacteria relative to non-diazotrophic cyanobacteria were used as the positive enrichment set; the remaining proteins in the 31,028-protein atlas universe were used as the comparator. Under the strictest primary-label exclusion, 237 diazotrophy-conserved proteins remained after removing condensate training positives and their homologous clusters.

Diazotrophy-conserved proteins had higher driver scores than the atlas background in the nitrogenase-withheld sensitivity analysis. Under the most conservative enrichment test, which removes nitrogenase-family labels from training, every condensate training positive, and every member of any MMseqs2 cluster containing a training positive, diazotrophy-conserved proteins showed a significant continuous score shift in the primary curated set (Mann-Whitney one-sided p = 1.66 × 10^-6). The same score-shift test remained significant in the conservative 23-positive set (p = 5.85 × 10^-14) and in the homolog-expanded 53-positive set (p = 3.23 × 10^-17). A seed-42 diagnostic effect-size calculation using the deposited family mapping gives common-language AUC 0.589 and rank-biserial correlation 0.178 for driver scores, consistent with a modest continuous shift rather than a large fixed-threshold effect.

The prespecified odds-ratio-at-threshold analysis was more sensitive to label-set composition. At driver-score >= 0.5 under the strict cluster exclusion, the conservative set gave OR 3.43 [1.44, 6.02], whereas the primary curated set gave OR 0.72 [0.14, 1.72] and the homolog-expanded set gave OR 0.87 [0.27, 2.41]. We therefore treat the continuous rank-sum result as the primary diazotrophy signal and the threshold odds ratios as sensitivity analyses. The nitrogenase-withheld model also changes the interpretation of Nif-family ranking. Reported ranks are computed on the 31,021-protein cyanobacterial sub-universe. In the primary curated set, current-study NifH1 (P00457) ranked at median 1,497 and current-study NifH2 (O30577) ranked at median 4,429 under the driver model, corresponding to the highest-scoring 4.8% and 14.3% of cyanobacterial proteins, respectively. NifK ranked at median 1,827 (highest-scoring 5.9%), NifB at 2,992 (highest-scoring 9.6%), NifE at 12,375, and NifN at 21,453. The NifH-GFP imaging result (Section 3.2) is therefore the direct biological observation, while the atlas supplies a broader prioritization and enrichment framework.

Under the same nitrogenase-withheld condition, feature ablation suggests that residual Nif-family signal is not carried by a single feature class. At seed 42, sequence-only ranks placed P00457 and O30577 at the 20.2% and 23.6% cyano-rank percentiles, whereas ESM-only ranks placed them at 40.5% and 53.2%. For NifK, the pattern was different, with ESM-only ranking strongest among the three feature sets at the 1.8% percentile. This mixed behavior argues against a simple pathway-memorization explanation and supports treating the atlas as a hypothesis-generation tool.

Beyond the nitrogenase core, the strongest primary shortlist signal is not a single metabolic pathway. It includes CcmM and other carboxysome-associated proteins, FtsZ/SepF-family cell-division proteins, SSB/RecA-family DNA maintenance proteins, RNA-binding proteins, chaperone/protein-quality-control proteins, cytochrome c6, and selected signaling or stress proteins. These results motivate, but do not prove, testing whether NifH-GFP puncta have a spatial relationship with thylakoid-associated reductant and ATP supply pathways.

The current computational and imaging data motivate a working model in which heterocyst-restricted NifH-GFP puncta could have a spatial relationship with thylakoid-associated reductant and ATP supply pathways. This remains a hypothesis: direct tests of focus position, endogenous nitrogenase composition, dynamics, ultrastructure, and local oxygen remain outstanding.

### 3.6 Wet-Lab Prioritization Shortlist

We generated a 100-protein prioritization shortlist from the primary curated nitrogenase-withheld cyano-only ranking (Supplementary Table S5). The driver-validation track is led by CcmM/carboxysome-associated proteins, FtsZ/SepF-family cell-division proteins, SSB and RecA DNA-maintenance proteins, Dps/ferritin stress proteins, RNA-binding proteins, GrpE/GroEL/DnaK-family chaperone systems, cytochrome c6, and selected signaling proteins. This shortlist is intended as a test queue, not a claim that each high-scoring protein forms a condensate.

A practical validation path would begin with purified-protein turbidity, microscopy, and concentration-dependent phase diagrams for candidate drivers. Candidate clients should then be tested for partitioning into NifH-GFP foci or other characterized bacterial condensates, with FRAP, 1,6-hexanediol sensitivity,crowding-agent controls, expression-level controls, and orthogonal localization tags before pathway-level claims are made.

## 4. Discussion

This study combines fluorescence microscopy of NifH-GFP in living heterocysts with a cyanobacterial sequence-based condensate-prioritization framework. The imaging experiment shows that NifH-GFP forms many discrete heterocyst-restricted puncta that are absent from constitutive GFP control and a heterocyst-expressed GFP comparator. The atlas provides a curated ranking resource and shows that diazotrophy-associated proteins remain shifted toward higher condensate-driver scores in nitrogenase-withheld sensitivity analyses.

The biological interpretation is narrower than a phase-separation claim. We describe the observed structures as NifH-GFP puncta in *Anabaena* sp. PCC 7120, not as proof that these foci are liquid condensates or oxygen-protective compartments. Heterocysts already combine multiple oxygen-protection strategies, including envelope specialization, downregulated photosystem II activity, and elevated respiration. The current data add a candidate subcellular organization pattern for nitrogenase, but FRAP, perturbation, composition, ultrastructure, and local oxygen measurements are needed before the foci can be assigned a mechanism.

### 4.1 Condensate-Mediated Organization of Cyanobacterial Metabolism

The atlas suggests that condensate propensity may be unevenly distributed across cyanobacterial systems. The deposited diazotrophy-conservation family table, literature-curated flux-balance analysis, and the sequence-based condensate atlas point to overlapping nitrogen-fixation and energy-metabolism questions, but these are computationally linked analyses rather than independent demonstrations of phase separation.

The primary shortlist is biologically coherent, drawing from CcmM and carboxysome-associated proteins as well as cell-division factors, DNA-maintenance proteins, RNA-binding proteins, chaperones, and electron-transfer proteins. That pattern fits broad bacterial condensate biology, but these data do not yet show co-localization, flux effects, or condensate formation for those proteins.

The electron-transfer signal is best read as biological motivation, not evidence for a PSI condensate mechanism. Photosystem I transfers electrons to ferredoxin, ferredoxin distributes reducing equivalents to multiple metabolic sinks, and ferredoxin-NADP+ reductase localization can influence linear versus cyclic electron flow^22-24^. Separately, pyrenoid biology shows that phase-separated photosynthetic carbon-fixation compartments can interface with thylakoid-associated electron-transfer machinery^25,26^. These precedents make the spatial relationship between NifH-GFP puncta and local reductant or ATP-supply pathways worth testing, but they do not show that ferredoxin organizes PSI condensates or that NifH-GFP puncta have a defined redox function.

### 4.2 Cross-Validation with Independent Computational Approaches

The nitrogen-fixation enrichment analysis (Section 3.5) provides an orthogonal consistency check using the deposited diazotrophy-conservation family mapping. In the primary curated atlas, proteins enriched for conservation across diazotrophic cyanobacteria score higher than background under the strictest sensitivity test after removing nitrogenase-family labels from training, all condensate training positives, and homologous clusters of those positives. This supports the idea that nitrogen-fixation biology is shifted toward higher condensate-propensity predictions, but it does not establish condensate formation for the diazotrophy-conserved proteins. The proportion of diazotrophy-conserved proteins above a fixed 0.5 score threshold was label-set sensitive and should be interpreted only as a sensitivity analysis.

The external LLPS-predictor comparison reinforces this interpretation. catGRANULE 2.0 and the atlas distinguish the curated benchmark labels at similar scales, yet their per-protein rankings are only modestly concordant, indicating overlapping but non-identical sequence signals. That partial independence makes the NifH result more informative: catGRANULE scores nitrogenase iron proteins highly, and the deliberately conservative nitrogenase-withheld atlas still ranks current-study NifH1 in the highest-scoring 4.8% and NifH2 in the highest-scoring 14.3% after removing nitrogenase-family labels and homologous clusters from the training data. Together with the continuous diazotrophy score shift across label sets and the feature-ablation result showing that both classical sequence features and PLM embeddings carry signal, these observations raise the priority of NifH-associated condensate biology without converting the imaging result into a mechanistic LLPS claim.

The ESM-2 embedding approach is not pathway-specific, so the workflow should transfer technically to other proteomes with appropriate labels and negatives. Biological generalization remains untested outside the seven cyanobacterial proteomes analyzed here.

### 4.3 Label Noise and Proxy Negatives

The ranking task inherently involves label noise: condensate-evidence Tier 2 proteins with punctate localization or microcompartment roles may not all be true LLPS drivers, and proxy negatives may include undiscovered condensate proteins. The low-IDR and phylo-matched negative strategies are useful sanity checks, but their near-perfect separation is likely inflated by negative-set construction; the random-proteome and homology-aware evaluations are the more conservative estimates of discriminative performance.

### 4.4 Limitations

Even after direct label restoration, the model has limited statistical power. The primary driver target contains 17 positives, and only two direct condensate-evidence Tier 1 driver anchors, SSB and FtsZ, are present. Both are heterologous bacterial proteins rather than cyanobacterial validation cases. The 27 associated positives also mix clients, scaffold-associated proteins, and microcompartment components. The feature-ablation result (Section 3.1) further shows that, under homology-aware CV, the 27 hand-crafted biophysical features perform comparably to ESM-2 650M embeddings for the driver target. Thus, the model is best presented as a joint biophysical-feature and PLM-embedding framework for ranking cyanobacterial condensate candidates, not as evidence that PLM embeddings alone reveal a new bacterial LLPS signal.

The wet-lab evidence is qualitative and rests on one NifH-GFP fusion in one organism, *Anabaena* sp. PCC 7120. Fluorescent proteins can perturb localization, oligomerization, and condensate behavior in tag- and fusion-context-dependent ways, including through weak GFP-family dimerization when non-monomeric variants are used^18-21^. The constitutive GFP control, strain A1198 (P*psbA1*-GFP), and the nifB-linked GFP reporter/comparator, strain A2094 (P*nifB*-*nifB*-GFP), argue against non-specific GFP aggregation as the only explanation (Figure 2b, d). However, A2094 is not a GFP-alone control, and we do not yet have a true heterocyst-specific GFP-alone control, monomeric or orthogonal tag variants, endogenous-locus tagging, FRAP-derived recovery half-times or mobile fractions, an isogenic PopZ-GFP positive control in the same strain, 1,6-hexanediol perturbation, or O_2_ biosensor data. The LLPS interpretation should therefore be treated as a working model anchored on one supportive observation with limited controls.

Only NifH has been tested at the bench in this work. Other top-ranked candidates, including CcmM/carboxysome-associated proteins, FtsZ/SepF, SSB, RecA, RNA-binding proteins, chaperones, cytochrome c6, and selected stress or signaling proteins, remain computational predictions. Local O_2_ was not measured, so the idea that NifH-GFP puncta occupy a locally reduced-O_2_ environment remains untested. The diazotrophy-conserved enrichment set comes from the deposited family-to-atlas mapping used for Figure 4, and the full high-confidence conservation-enriched family list and broader diazotrophy-associated family list are deposited with this manuscript for reproducibility.

**Figure 4.**
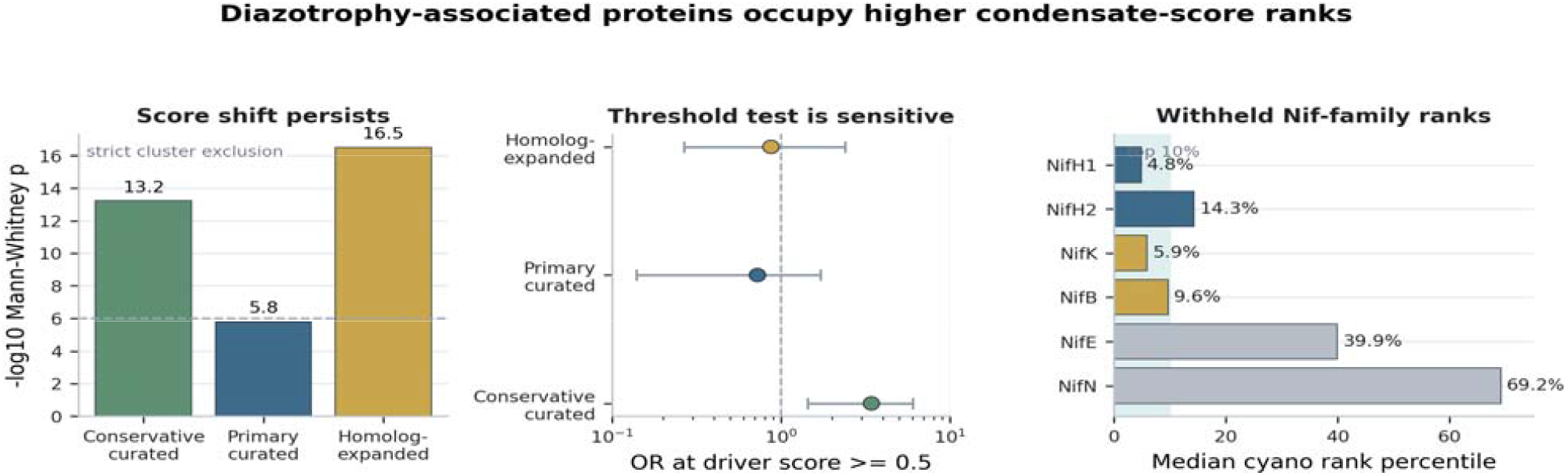
Diazotrophy-associated proteins occupy higher condensate-score ranks under nitrogenase-withheld sensitivity models. **(A)** Continuous score-shift evidence for proteins enriched for conservation across diazotrophic cyanobacteria under the strict cluster-exclusion test, shown as -log10 Mann-Whitney one-sided p values across conservative curated, primary curated, and homolog-expanded label sets. **(B)** Prespecified threshold test at driver-score >= 0.5, shown as bootstrap odds ratio and 95% interval; this threshold-form enrichment is label-set sensitive and is not used as the headline claim. **(C)** Median cyano-only rank percentiles for NifH/K/B/E/N-family proteins under the primary curated nitrogenase-withheld model.

**Figure 5.**
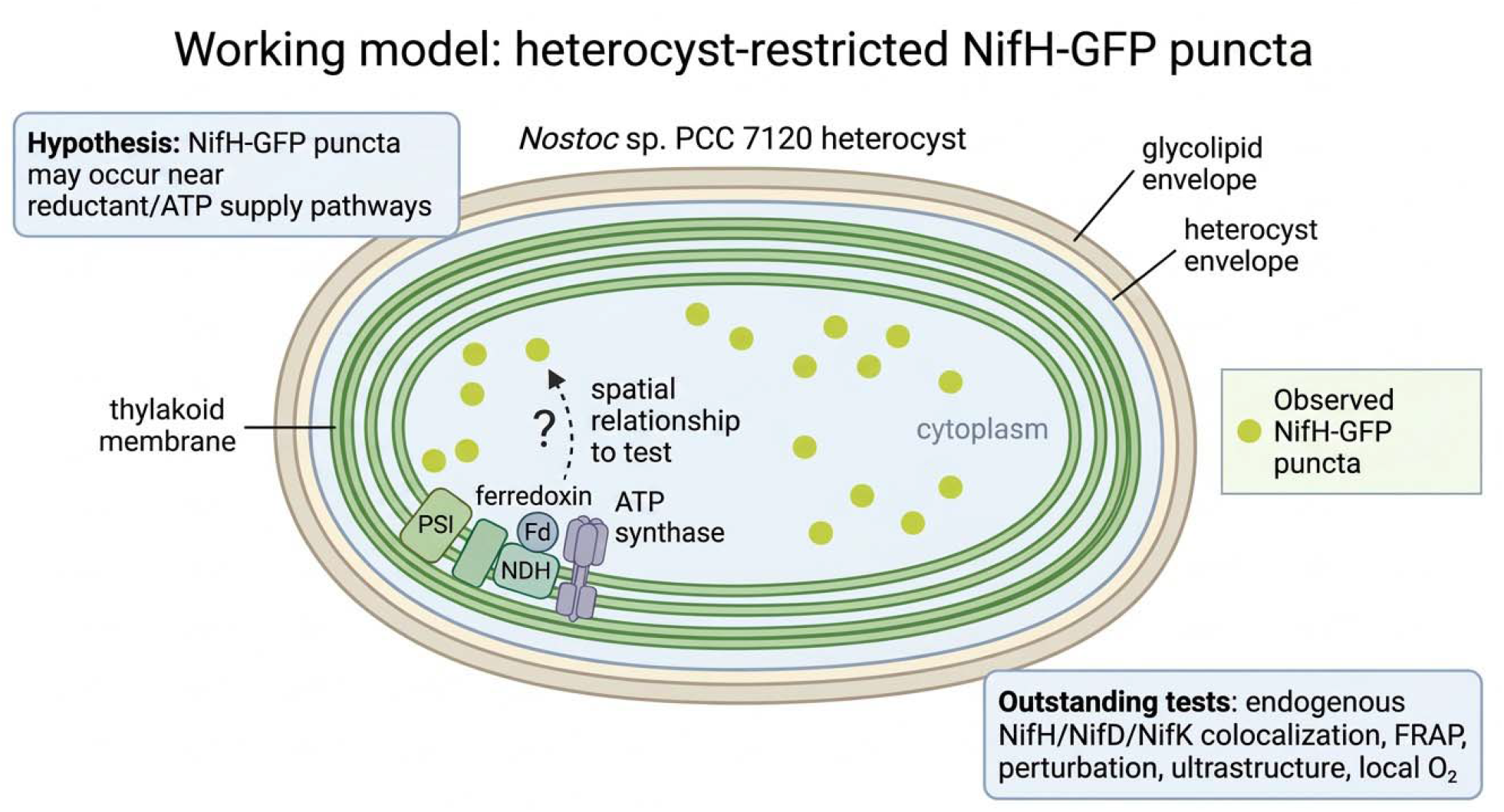
Working model: heterocyst-restricted NifH-GFP puncta. Cross-section of an *Anabaena* sp. PCC 7120 heterocyst showing observed NifH-GFP puncta and the hypothesis that their spatial relationship to thylakoid-associated reductant and ATP supply pathways should be tested. The glycolipid and heterocyst envelope layers are shown as known features of heterocyst biology. Endogenous NifH/NifD/NifK colocalization, FRAP, perturbation, ultrastructure, and local O_2_ measurements remain outstanding.

### 4.5 Future Directions

The next experimental step is a systematic test of the wet-lab prioritization shortlist (Section 3.6) using purified-protein turbidity, microscopy, concentration-dependent phase diagrams, FRAP, perturbation, and appropriate positive and negative controls. Future computational iterations should incorporate structural features where they add information, expand the positive-label set as more cyanobacterial condensate studies emerge, and test whether the same workflow transfers to other microbial lineages. Context-dependent condensate formation during heterocyst differentiation, nitrogen fixation induction, and stress responses remains an especially important target for future work. Condensates can be gated by concentration, multivalency, pH, ionic strength, energy state, stress, and the surrounding cellular environment^1,27-29^. The NifH-GFP phenotype may therefore reflect a state-dependent assembly in which heterocyst maturation, nitrogenase expression, redox demand, ATP demand, and oxygen-management physiology provide the context that permits NifH-associated foci to form.

## 5. Methods Availability

Processed data tables, figure source files, and analysis scripts are available from the corresponding author upon reasonable request and will be deposited in a public repository before journal submission. config/project_spec.yaml specifies the main parameters, and scripts 01-10 plus the rescue and sensitivity scripts reproduce the unified workflow. Corrected labels, label-audit tables, three-label-set sensitivity comparisons, nitrogenase-withheld rankings, cross-validation metrics, enrichment summaries, external-benchmark outputs, and wet-lab shortlist files are provided under data/curated/ and results/tables/. Strict-audit and rescue provenance files are provided under docs/hpc_review_20260509/, docs/hpc_rescue_20260509/, logs/, and scripts/. The requirements.txt file records package versions where available, and tests/test_minimal.py covers the two core utilities (compute_sequence_features, compute_metrics). All numerical results can be regenerated from data/raw/ onward; local feature generation takes about 1 hour on an RTX 3070 GPU, and sensitivity reruns were performed on SDSU HPC using the same processed feature assets. The external-predictor benchmark (catGRANULE 2.0, PSAP, LLPhyScore) reported in Section 3.1 was run on a Linux HPC (RHEL 9, Python 3.9); per-tool install recipes, raw scores, and the benchmark manifest are provided at results/tables/external_benchmark/, and the SOP for reproducing the run is in docs/HPC_EXTERNAL_BENCHMARK_RUNBOOK.md.

## Supporting information

Nitrosome Video

## Acknowledgments

This research was supported in part by South Dakota State University’s High Performance Computing resources.

## Supplementary Figures

**Supplementary Figure S1.**
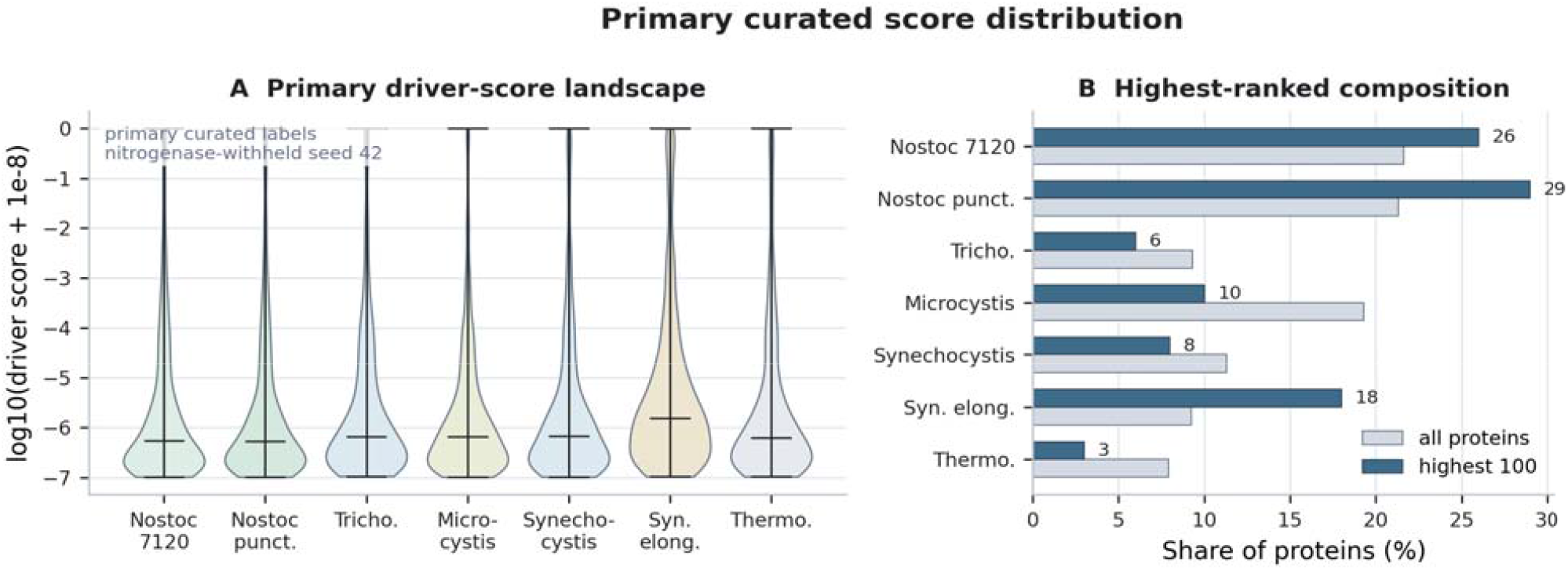
Primary curated nitrogenase-withheld driver-score distributions.**(A)** Driver-score distributions across the seven cyanobacterial proteomes for the seed-42 cyano-only ranking, plotted as log10(score + 1e-8). **(B)** Organism composition of all proteins compared with the highest-ranked 100 non-training, non-nitrogenase-withheld candidates. The diazotrophy analysis in Figure 4 operates within the atlas universe rather than as a simple species-morphology effect.

**Supplementary Figure S2.**
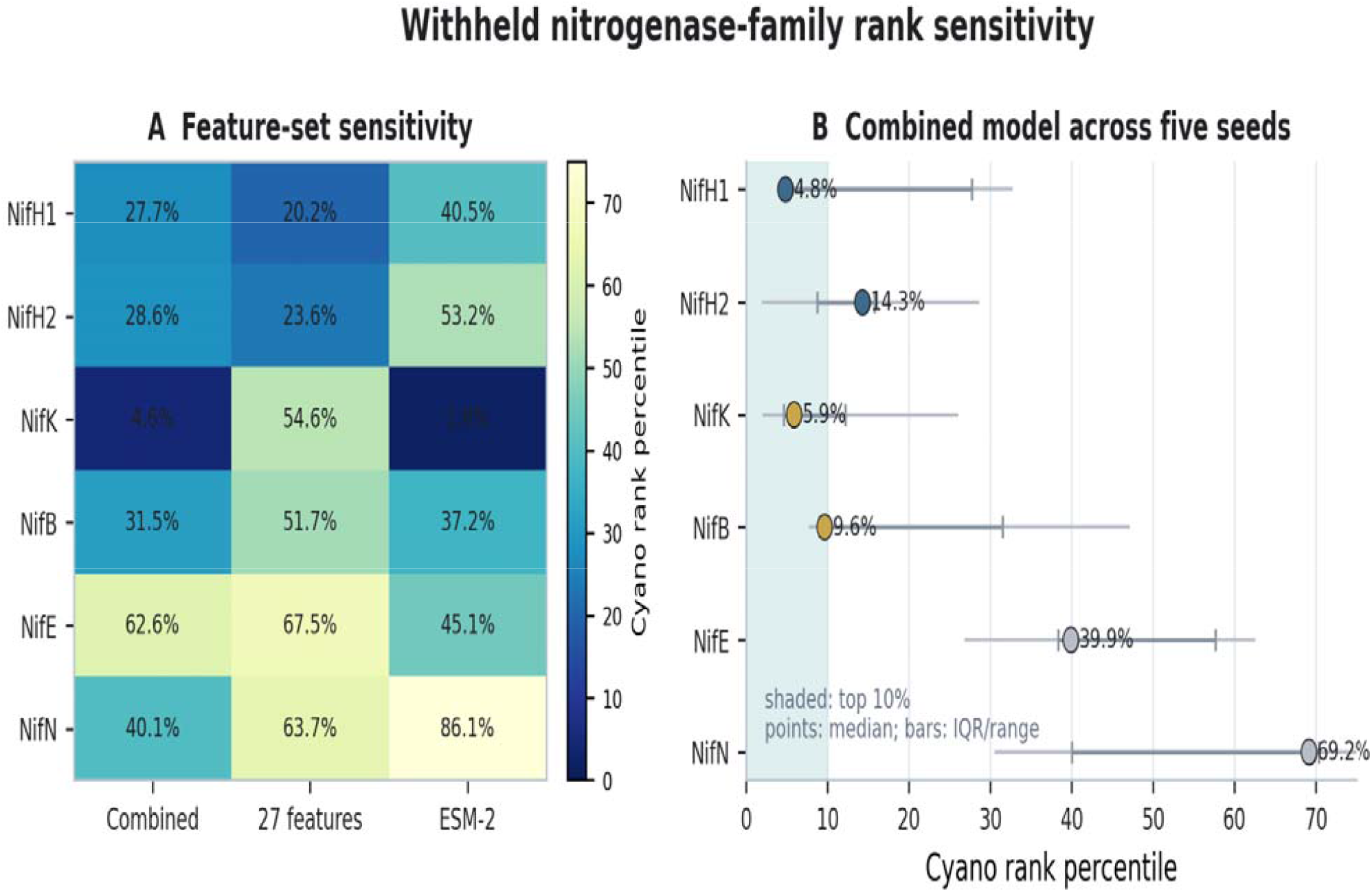
Feature-set and seed sensitivity of nitrogenase-family driver ranks under the primary curated nitrogenase-withheld model.**(A)** Cell values show seed-42 cyano-only rank percentiles for selected NifH/K/B/E/N-family proteins under the combined, sequence-feature-only, and ESM-only models; lower percentiles indicate stronger ranks. **(B)** Five-seed combined-model rank percentiles are shown as medians with interquartile ranges and full ranges. The strongest feature set differs by protein, supporting use of the atlas as a prioritization tool rather than a single-feature mechanistic classifier.

**Supplementary Figure S3.**
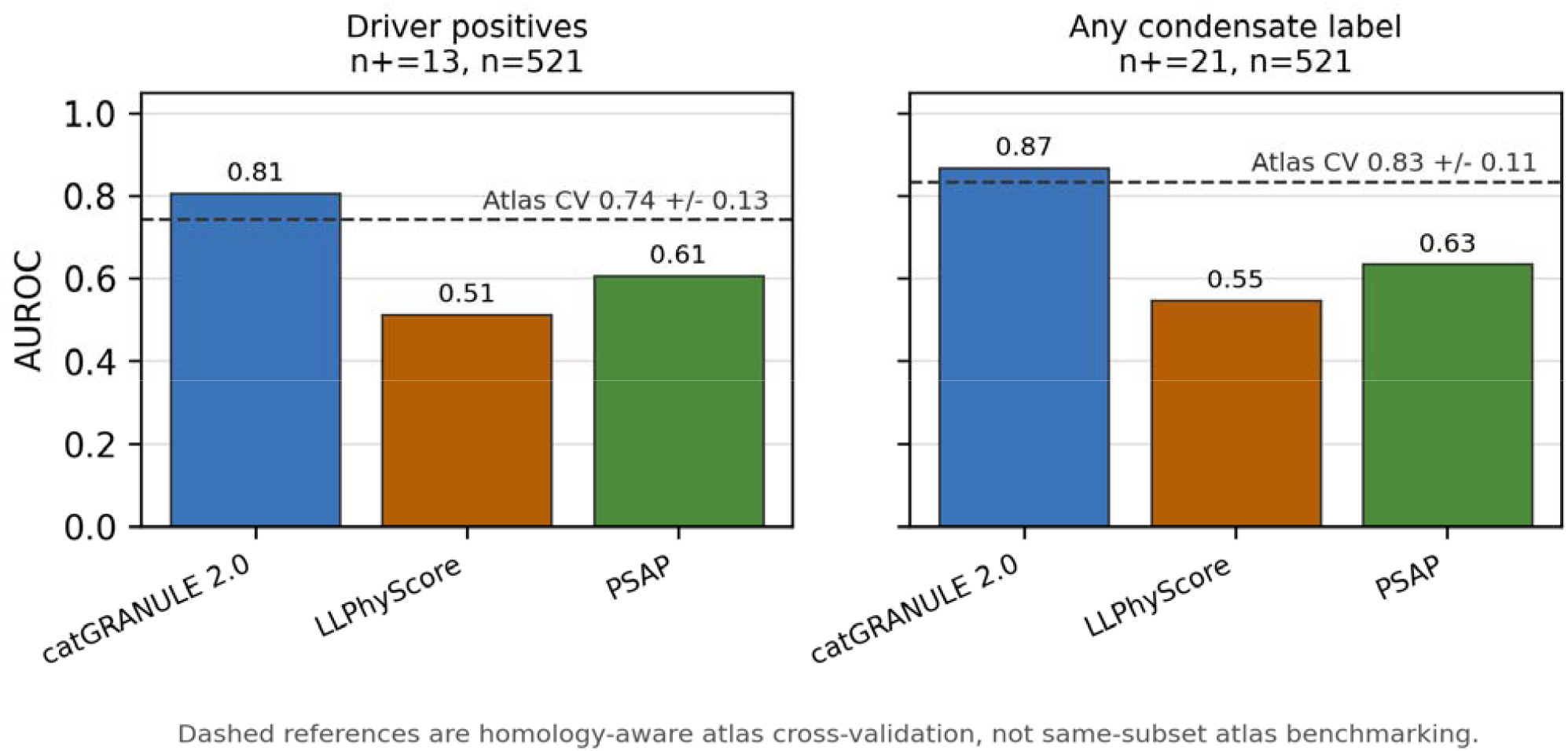
External LLPS-predictor benchmark on the conservative cleaned labeled-plus-500-cyano-background subset retained from the HPC audit. The subset contains 13 driver positives, 21 any-condensate positives, and 500 background proteins. catGRANULE 2.0, PSAP, and LLPhyScore were evaluated on Linux HPC; atlas homology-aware CV AUROCs are shown only as dashed reference lines because same-subset atlas benchmark AUROCs overlap training labels. The dashed atlas references are the conservative 23-positive CV values; the primary curated 44-positive CV values are 0.827 +/-0.142 for driver labels and 0.829 +/-0.096 for any-condensate labels. catGRANULE 2.0 achieved AUROC 0.81 on driver labels and 0.87 on any-condensate labels, while its per-protein ranks were only modestly concordant with atlas ranks, supporting complementary rather than redundant prioritization. PSAP achieved 0.61 and 0.63. LLPhyScore achieved 0.51 on driver labels and 0.55 for any-condensate labels after score-orientation flipping; its published score direction for any-condensate labels was below chance, which may also be partially attributed to low sample size.

**Supplementary Figure S4.**
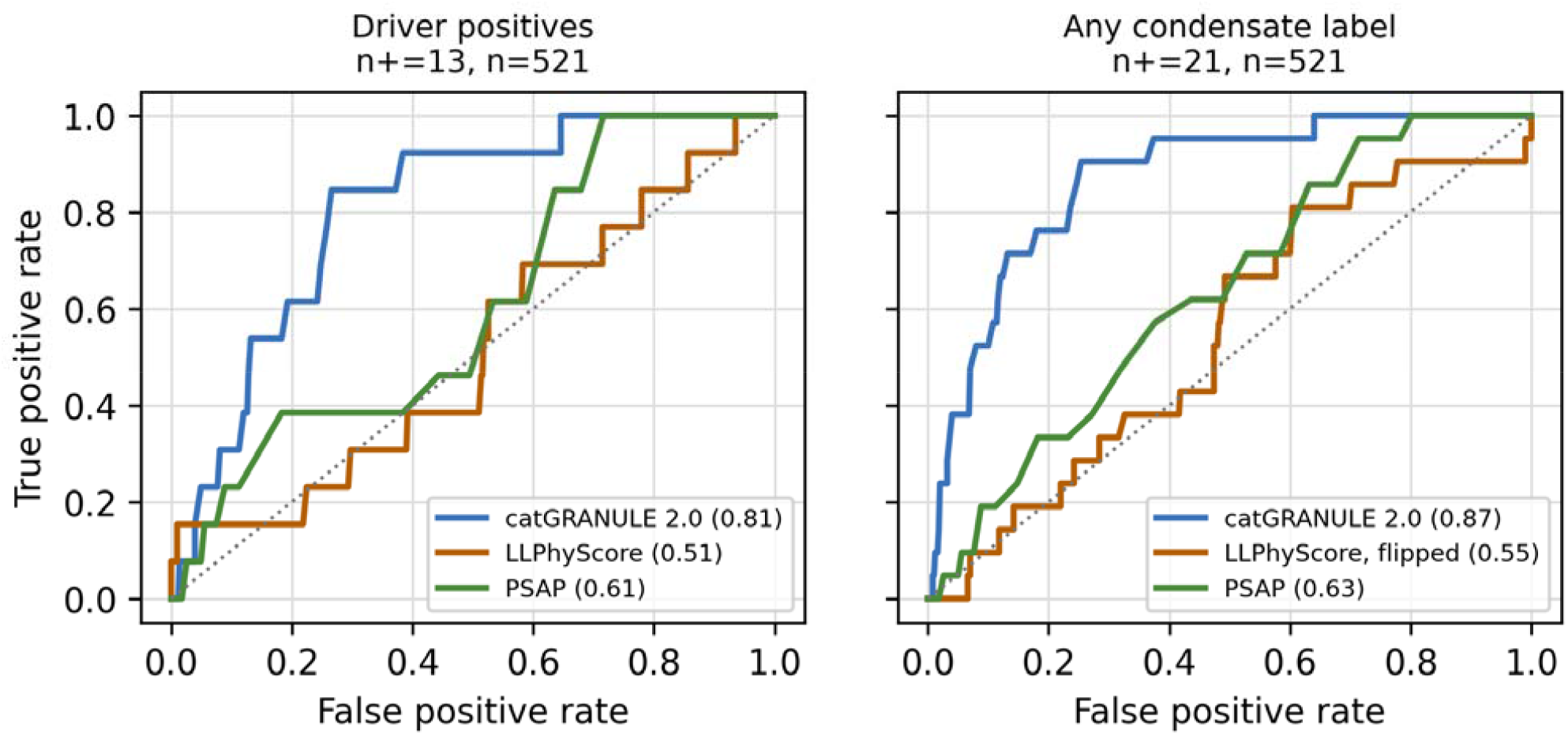
Receiver-operating-characteristic (ROC) curves for three eukaryote-trained external LLPS predictors on the conservative 521-protein labeled-plus-500 benchmark subset used for the external-predictor comparison. **(A)** Driver labels (n = 13 positives). **(B)** Any-condensate labels (n = 21 positives). catGRANULE 2.0 is the strongest external predictor (AUROC 0.81 driver / 0.87 any-condensate); PSAP reaches 0.61 / 0.63; LLPhyScore reaches 0.51 / 0.55 when the any-condensate score direction is flipped. The atlas same-subset in-sample ROC curve is intentionally omitted because it is not a generalization estimate. The dashed gray line is chance. Per-tool scores are in results/tables/external_benchmark/, and the conservative benchmark summary is in results/tables/sensitivity/external_benchmark_hpc_review.json.

**Supplementary Figure S5.**
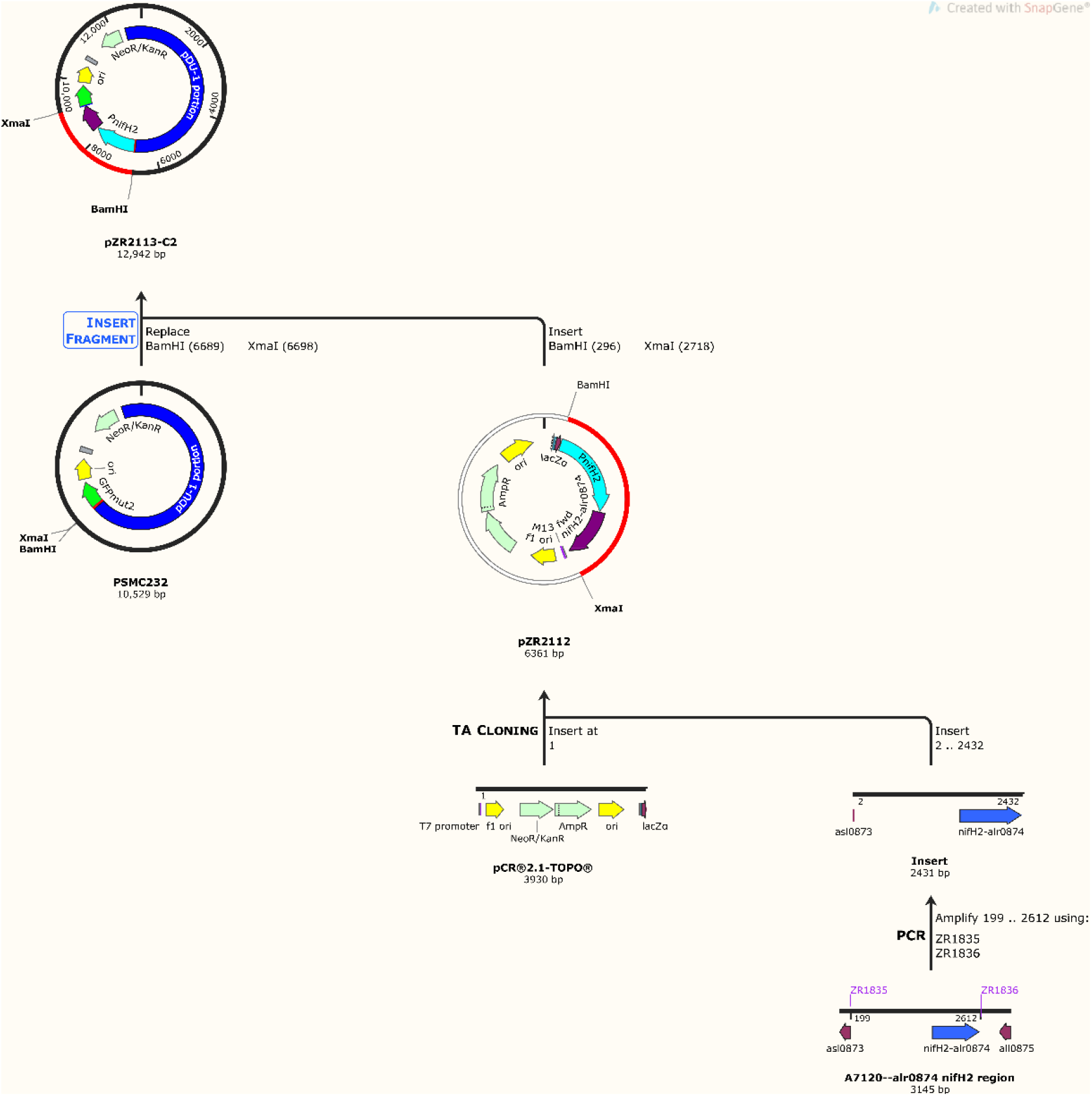
Construction of pZR2113-C2 (P*nifH2*-NifH-GFP). The 12,942-bp shuttle plasmid was assembled by inserting a PCR-amplified *Anabaena* 7120 *nifH2*–*alr0874* region (primers ZR1385 / ZR1386) into pCR®2.1-TOPO® via TA cloning to give intermediate pZR2112, then sub-cloning the BamHI– XmaI fragment of pZR2112 into the pDU1-derived shuttle vector pSMC232 to yield pZR2113-C2. Details in Tables S1 and S2.

**Supplementary Figure S6.**
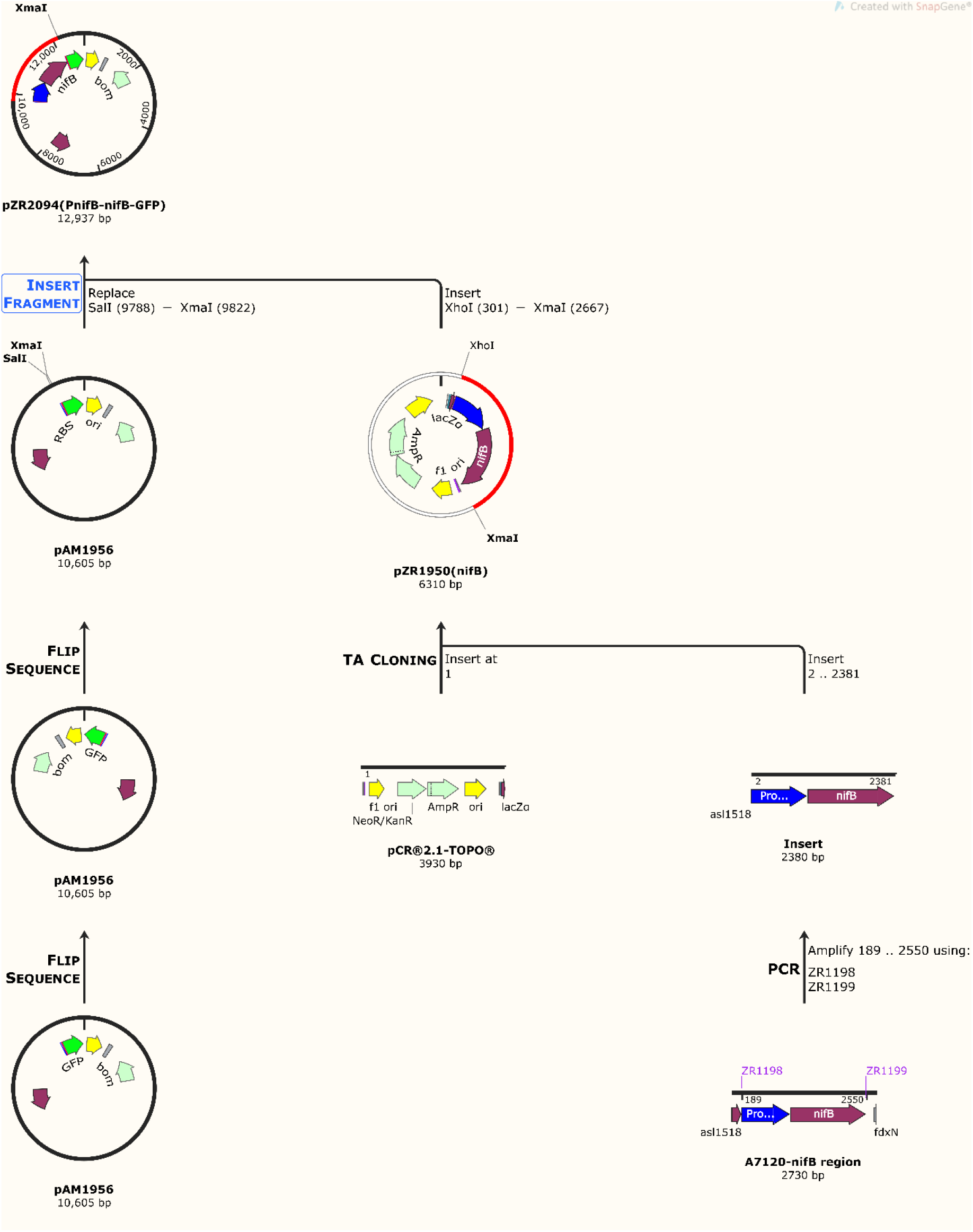
Construction of pZR2094 (P*nifB*-*nifB*-GFP), the nifB-linked GFP reporter/comparator plasmid. The 12,937-bp shuttle plasmid was assembled by inserting a PCR-amplified *Anabaena* 7120 *nifB* promoter–*nifB* region (primers ZR1198 / ZR1199) into pCR®2.1-TOPO® via TA cloning to give intermediate pZR1950(*nifB*), then sub-cloning the XhoI–XmaI fragment into a flipped-orientation derivative of pAM1956 to yield pZR2094. Details in Tables S1 and S2.

**Supplementary Figure S7.**
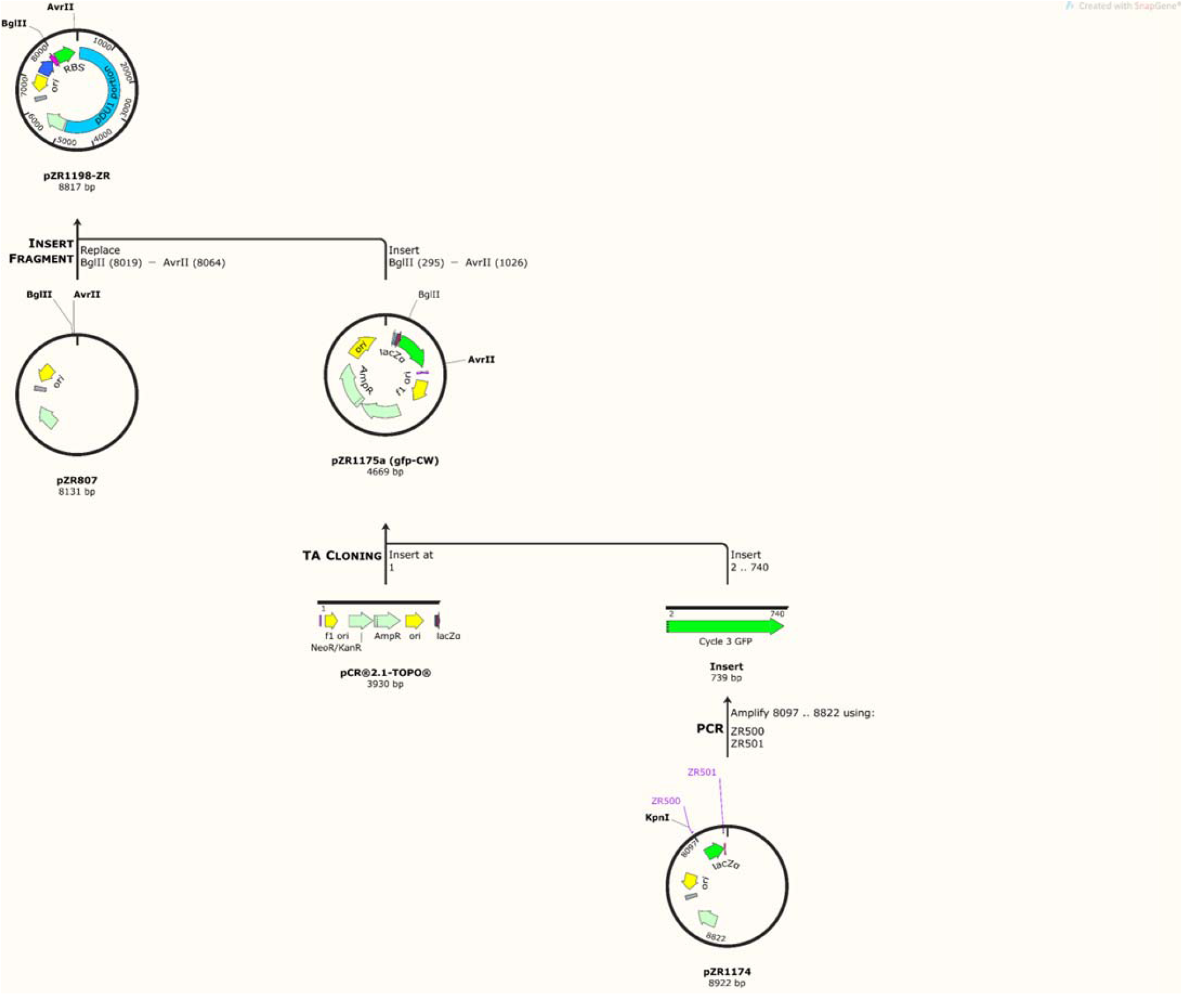
Construction of pZR1198 (P*psbA1*-GFP), the constitutive GFP-only control plasmid. The 8,817-bp shuttle plasmid was assembled by inserting a PCR-amplified *gfp* coding region (primers ZR500 / ZR501) into pCR®2.1-TOPO® via TA cloning to give pZR1175a, then sub-cloning the BglII– AvrII fragment into pZR807 (P*psbA1* shuttle backbone) to yield pZR1198. Details in Tables S1 and S2.

**Supplementary Figure S8.**
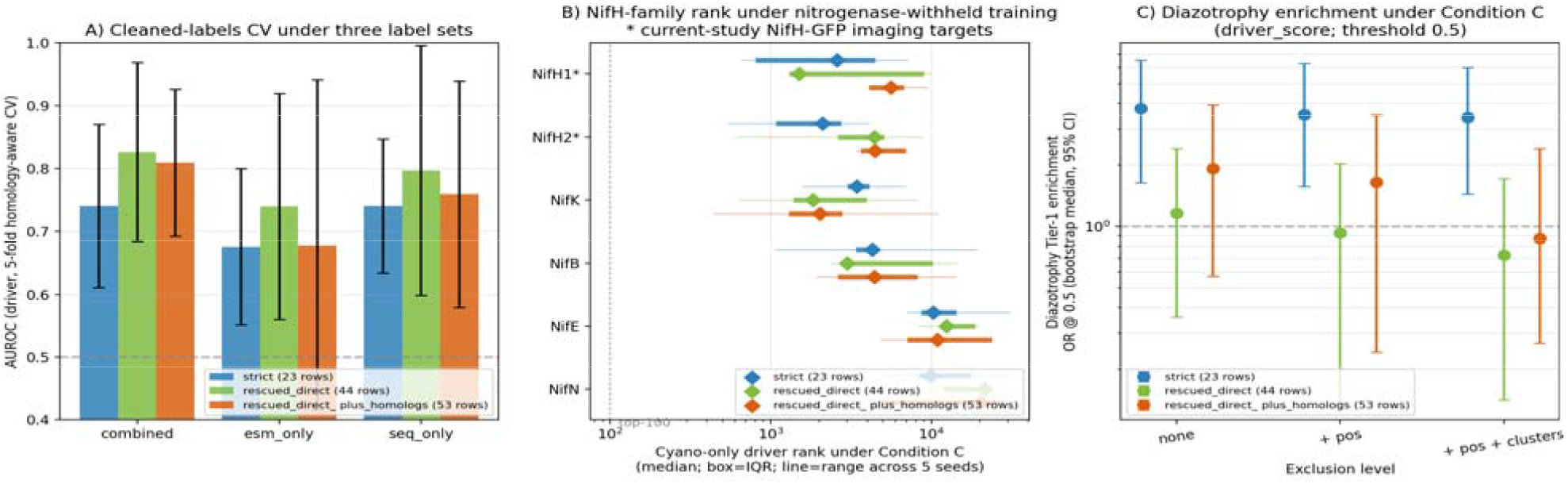
Three-label-set sensitivity analysis comparing the conservative curated, primary curated, and homolog-expanded label sets. The figure summarizes driver AUROC, nitrogenase-withheld NifH rank behavior, and diazotrophy threshold sensitivity. These results support using the primary curated set for manuscript claims while treating fixed-threshold odds ratios as sensitivity analyses rather than headline enrichment results.

## Supplementary Movie

**Movie S1**. Three-dimensional z-stack reconstruction of one mature heterocyst in *Anabaena* sp. PCC 7120 strain A2113-C2 expressing NifH-GFP, rotated through the optical axis to reveal the spatial distribution of discrete NifH-GFP puncta filling the heterocyst volume. Confocal imaging through the GFP filter; a representative z-stack reconstruction is shown in Figure 2c.

## Supplementary Tables

**Table S1**. Plasmids and bacterial strains are used in this study.

**Table S2**. Oligonucleotide primers are used for plasmid construction.

**Table S3**. Prespecified fixed-threshold diazotrophy-enrichment sensitivity summary. Bootstrap odds-ratio medians, 95% intervals, raw Fisher exact p values, and within-label-set Benjamini-Hochberg adjusted p values for driver-score thresholds 0.5, 0.7, and 0.9 are provided in results/tables/sensitivity/threshold_enrichment_summary_supplement.tsv. Continuous Mann-Whitney U tests remain the primary diazotrophy score-shift evidence.

**Table S4**. Diazotrophy-conserved family-to-atlas mapping used for Figure 4. The table is provided in results/tables/tier1_condensate_joined.csv; the seed-42 diagnostic effect-size summary is provided in results/tables/sensitivity/diazotrophy_score_shift_effect_size_seed42.tsv.

**Table S5**. Wet-lab prioritization shortlist. The 100-protein shortlist from the primary curated nitrogenase-withheld cyano-only ranking is provided in results/tables/wetlab_shortlist_unified.tsv.

**Table S6**. External-predictor concordance and diagnostic matched-null outputs. Per-tool scores are provided in results/tables/external_benchmark/; the seed-42 length- and IDR-matched nitrogenase-family diagnostic null is provided in results/tables/sensitivity/nif_family_length_idr_matched_null_seed42.tsv.

**Table S1.** Plasmids and Bacterial Strains Used in This Study.

| **Plasmids** |  |  |  |
| --- | --- | --- | --- |
| **Name** | **Characteristics** | **Construction** | **Source** |
| pSMC232 | Km<sup>R</sup>/Nm<sup>R</sup>; MCS;<br>SRGSASAS-gfpmut2;<br>promoterless gfpmut2<br>translational fusion vector |  | [5] |
| pAM1956 | Nm<sup>R</sup>, transcriptional fusion<br>GFPmut2-containing cloning<br>vector |  | [6] |
| pRL443 | Ap<sup>R</sup>, Tc<sup>R</sup>; Conjugal Plasmid |  | [9] |
| pRL623 | Cm<sup>R</sup>; Helper Plasmid |  | [9] |
| pCR2.1-TOPO | Km<sup>R</sup>/Ap<sup>R</sup>; TOPO-TA cloning<br>vector |  | Invitrogen |
| pZR807 | Km<sup>R</sup>; |  | [10] |
| pZR606 | Km<sup>R</sup>/SP<sup>R</sup>; integration vector |  | [7] |
| pZR666 | MCS-uvgfp-MCS cassette; | LK2406/LK2407 annealed oligo ligated<br>into AflII digested pZR606 | This study |
| pZR1158b | Km<sup>R</sup>/Ap<sup>R</sup>; | Primers ZR481, ZR482 to PCR amplified<br>uv-gfp from pZR666 and ligated to<br>pTOPO2.1 to produce pZR1158b | This study |
| pZR1174 | Km<sup>R</sup>; | KpnI + SpeI to get uvgfp from pZR1158b-5<br>and ligated to KpnI + AvrII digested<br>pZR1157-8 to produce pZR1174. |  |
| pZR1157 | Km<sup>R</sup>; | Annealed oligo ZR479/480 ligated to BglII<br>+ KpnI digested pZR807-7 to produce<br>pZR1157. | This study |
| pZR1175a | Km<sup>R</sup>/Ap<sup>R</sup>; | Primers ZR500, ZR501 to PCR amplify uv-<br>gfp from pZR1174-1 and ligated the 733bp<br>gfp to pTOPO2.1 to produce pZR1175a |  |
| pZR1198 | Km<sup>R</sup>/Nm<sup>R</sup>; P<sub>nir</sub>-P<sub>psbA1</sub>-uvGFP | Previously developed in the Zhou Lab (Fig<br>S3 and details in Table S1 &S2) | This study |
| pZR1950 | Km<sup>R</sup>/Ap<sup>R</sup>; | Primers ZR1198/ZR1199 for PCR BamHI-XhoI-P<sub>nifB</sub>-nifB-TCC CGG G 2378bp using<br>A7120 genomic DNA and ligated to<br>pTOPO2.1 to produce pZR1950 (JG) | This study |
| pZR2094 | Km<sup>R</sup>/Nm<sup>R</sup>; P<sub>nifB</sub>-nifB-GFP | XhoI-P<sub>nifB</sub>-nifB-XmaI from pZR1950<br>ligated to SalI-XmaI digested pAM1956 to<br>produce pZR2094 (P<sub>nifB</sub>-nifB-GFP produces<br>two proteins, NifB and GFP because nifB-gfp was in transcriptional fusion). | This study |
| pZR2112 | Km<sup>R</sup>/Ap<sup>R</sup>; P<sub>nifH</sub>-NifH-GFP<br>cloned into pCR2.1-TOPO | Phusion High-Fidelity DNA Polymerase<br>PCR amplified BamHI/XhoI-<br>P<sub>alr0874</sub>(nifH2)-alr0874-TCC CGG G-2431bp using Anabaena 7120 genomic<br>DNA with PCR primers ZR1385 and<br>ZR1386 and then cloned the 2.4Kb<br>fragment to pTOPO2.1 to produce<br>pZR2112 | This study |
| pZR2113-C2 | Km<sup>R</sup>/Nm<sup>R</sup>; P<sub>nifH</sub>-NifH-GFP | BamHI-2431bp-XmaI from pZR2112 to BamHI-XmaI digested pSMC232 to produce pZR2113-C2, finally transfer into A7120 for expression of NifH2-GFPmut2 chimeric protein | This study |

Bacterial Strains
| Name | Description | Source |
| --- | --- | --- |
| HB101 | *E. coli* used for the conjugation procedure with *Anabaena* 7120 | This study |
| TOP10 | *E. coli* cloning host | Invitrogen |
| NEB10β | *E. coli* cloning host | New England Biolabs |
| *Anabaena* sp. PCC 7120 | *Anabaena* sp. PCC 7120 wildtype strain | This study |
| A2113-C2 | *Anabaena* sp. PCC 7120 harboring plasmid pZR2113 | This study |
| A1198 | *Anabaena* sp. PCC 7120 harboring plasmid pZR1198 | ZR Lab # 725 |
| A2094 | *Anabaena* sp. PCC 7120 harboring plasmid pZR2094 | ZR Lab # 4325 |

Acronyms: Ap<sup>R</sup> -ampicillin resistance; Cm<sup>R</sup>/Em<sup>R</sup> -chloramphenicol/erythromycin resistance; Nm<sup>R</sup>/Km<sup>R</sup> - neomycin/kanamycin resistance; Tc<sup>R</sup> -tetracycline resistance; MCS -Multiple Cloning Site

**Table S2.**
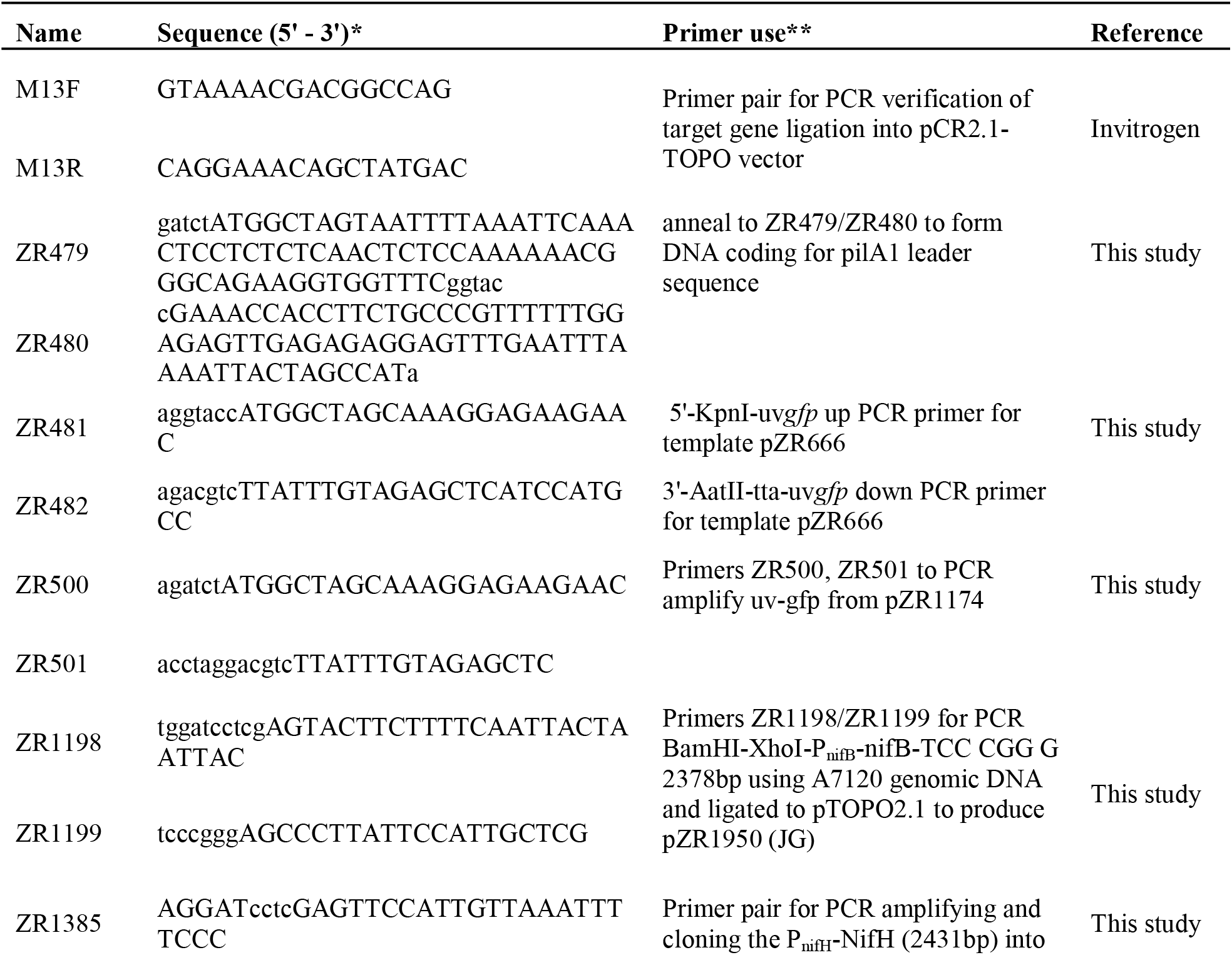
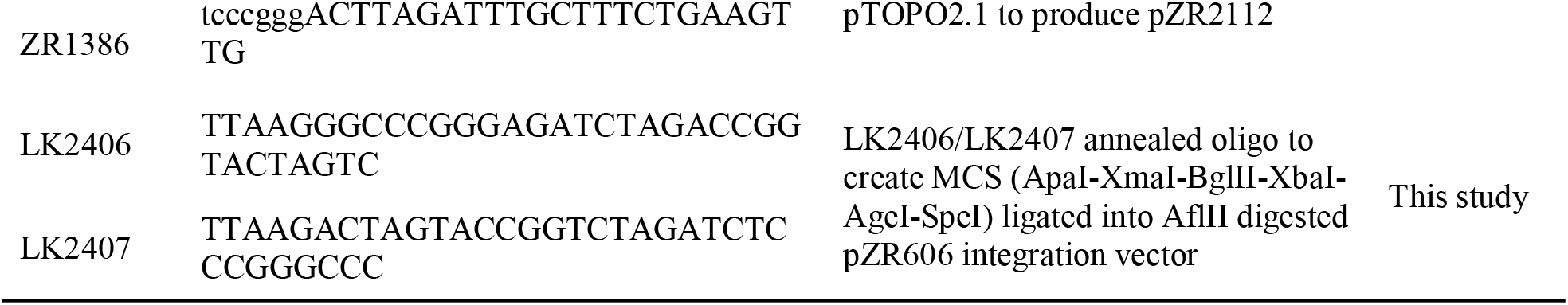
Primers Used in This Study.

